# PertDiffBench: Benchmarking Diffusion Models for Single-Cell Perturbation Response Prediction

**DOI:** 10.64898/2026.06.13.732013

**Authors:** Zijun Song, Yujia Xiang, Zhi Song, Wenyi Jin, Jiangmeng Li, Chuxiong Sun, Linhai Xie

**Author notes:** Corresponding author* **Emails:** Linhai Xie. Equal contributions (Zijun Song, Yujia Xiang).

## Abstract

Diffusion models are increasingly used to predict transcriptional responses to perturbations, but whether they improve on simpler generative and representation-based baselines remains unclear. Existing evaluations often do not separate the effects of model architecture, input representation, biological context and metric choice, making it difficult to determine where diffusion-based methods are useful. Here we introduce PertDiffBench, a standardized benchmark for diffusion-based transcriptomic perturbation prediction across single-cell and bulk RNA-seq datasets. PertDiffBench evaluates diffusion-based models across three complementary evaluation settings: standard prediction in known single-cell contexts and bulk perturbation conditions, generalization to unseen cell types, species, drugs and intermediate time points, and stress tests of feature dimensionality, input representation, noise type and gene ordering. Across these settings, diffusion models did not show a consistent advantage. scGen remained a strong baseline in common prediction tasks, whereas scDiffusion was the most competitive diffusion-based method in several generalization settings. Temporal imputation showed a different pattern, with a simple DDPM operating directly in expression space outperforming more specialized models. Stress tests showed that performance was model dependent and sensitive to feature dimensionality, encoder choice, noise type and gene ordering. Pretrained encoders did not consistently improve performance, with the classical scVI representation slightly exceeding STATE in seen-condition and unseen-cell-type settings. These results indicate that diffusion-model performance in perturbation response prediction depends strongly on task design and representation choice. PertDiffBench provides a practical framework for evaluating these models under biologically varied and stress-tested conditions.

## Introduction

Single-cell perturbation technologies have made it possible to study gene function and cell-state transitions at high resolution (Bunne, et al., 2024; Peidli, et al., 2024). By measuring transcriptional responses to genetic or chemical perturbations in individual cells, these assays can reveal state-specific effects, population heterogeneity and rare responsive subpopulations that are not accessible from bulk readouts. Yet the experimental space grows quickly with perturbation type, cellular context, dose, time and species, and soon exceeds what can be measured systematically. Given the cost, throughput and applicability limits of current single-cell assays, exhaustive experimental coverage is not feasible (Srivatsan, et al., 2020; Yao, et al., 2024). Predicting cellular responses to unmeasured perturbations and unseen biological conditions from limited observations has therefore become a central goal of single-cell perturbation modelling (Qian, et al., 2025; Rood, et al., 2024).

Computational methods have evolved with this goal, moving from the description of measured perturbation effects toward generalization across unmeasured conditions. Early methods such as scGen used variational autoencoders to model perturbation-induced shifts in latent space and predict responses in unmeasured cellular contexts (Lotfollahi, et al., 2019). CPA then extended this framework through compositional modeling of perturbation, dose and covariate effects, enabling predictions across doses, combinations, time points and species (Lotfollahi, et al., 2023). More recent knowledge-informed graph models and single-cell foundation models, including GEARS, scGPT, scFoundation, STATE and Tahoe-x1, have further broadened the scope of out-of-distribution perturbation response prediction (Adduri, et al., 2025; Cui, et al., 2024; Gandhi, et al., 2025; Hao, et al., 2024; Roohani, et al., 2024).

Diffusion models have recently entered this landscape as a new class of generative models for cellular response prediction. By learning conditional distributions through iterative denoising, they provide a flexible way to model heterogeneous and potentially multimodal cellular responses. Their formulation also allows perturbation identity, dose, cell state and other covariates to be included in the generation process. Early single-cell applications, including scVAEDer, scDiff and scDiffusion, explored diffusion-based generation of cellular states under specified biological conditions, with scVAEDer among the first to demonstrate perturbation response prediction using a VAE–diffusion framework (Luo, et al., 2024; Sadria and Layton, 2025; Tang, et al., 2023). More recent methods have become increasingly specialized: dbDiffusion focuses on unseen genetic perturbations and connects generation with debiasing and downstream statistical inference (Shang, et al., 2025). scPPDM addresses single-cell drug response prediction, with an emphasis on unseen drugs and covariate combinations (Liang, et al., 2025). Squidiff extends diffusion-based modelling to developmental dynamics and mechanistic interpretation (He, et al., 2025). In parallel, morphology-focused models such as MorphoDiff and MorphDiff apply diffusion models to perturbation-induced cellular imaging phenotypes (Navidi, et al., 2025; Wang, et al., 2025). These studies suggest that diffusion-based perturbation response prediction is emerging as a diverse family of methods with different architectures, conditioning strategies, modalities and prediction targets.

However, it remains unclear what actually drives the performance of diffusion models in omics perturbation prediction. Diffusion models were first successful in domains such as imaging, where nearby features show strong local dependence and the data have an explicit spatial structure. Omics data differ in an important way: genes have no natural spatial ordering, and their dependencies are often nonlocal, context dependent and shaped by regulatory programs rather than local neighbourhoods. This raises a central question for perturbation modelling. When diffusion-based methods improve performance, do the gains come from the diffusion backbone itself, or from other parts of the pipeline, such as input representation, feature selection, dataset scale, noise structure or task design?

Current benchmark studies do not yet answer this question. Existing comparisons differ in datasets, train/test splits, evaluation metrics and baseline selection, making results difficult to align across studies (Csendes, et al., 2025; Li, et al., 2025; Li, et al., 2025; Viñas Torné, et al., 2025; Wei, et al., 2026; Wenteler, et al., 2025; Wu, et al., 2024). Diffusion-based models have also not been evaluated systematically within a single framework, so their relative strengths and limitations remain difficult to assess. This gap is important because recent benchmarks suggest that complex models are not consistently superior to simpler baselines and that model rankings can vary across tasks and metrics. A unified benchmark is therefore needed to determine when diffusion models offer meaningful gains and which components of the modelling pipeline drive their performance.

Here we present ***PertDiffBench***, a benchmark for evaluating diffusion models in single-cell perturbation response prediction. PertDiffBench compares diffusion-based methods, including scDiffusion, scDiff, Squidiff and DDPM-based baselines, with established generative baselines such as scGen within a unified framework covering seen-condition prediction, unseen cell-type prediction, cross-species transfer, unseen mechanism-of-action prediction and temporal imputation. It also tests how performance is affected by feature dimensionality, data representation, input noise and feature ordering. By evaluating these factors in the same framework, PertDiffBench helps distinguish the contribution of the diffusion backbone from that of representation, data preprocessing and task design, and provides a clearer view of when diffusion models are useful for single-cell perturbation response prediction.

## Results

### Overview of PertDiffBench

We construct PertDiffBench to assess how well diffusion models support single-cell perturbation response prediction (Figure 1). The framework is designed to test whether performance gains are driven by the diffusion backbone itself or from other factors, including the input data representation, data scale, noise structure and the difficulty of the extrapolation task. We selected models that represent distinct conditional generation strategies in current single-cell perturbation modeling: scDiffusion uses classifier-guided conditional generation, scDiff incorporates conditions through a cross-attention-based diffusion framework, and Squidiff integrates semantic features into the denoising process to model cell-state transitions and perturbation responses. To provide controlled references, we further implemented two DDPM baselines: one that generates directly in the original gene-expression space and one that generates in a 256-dimensional latent space encoded by a three-layer MLP. We also included scGen as a classical VAE-based baseline for perturbation response prediction. This model panel allowed us to compare published diffusion architectures with simplified diffusion baselines and an established non-diffusion generative model under the same evaluation framework.

**Figure 1.**
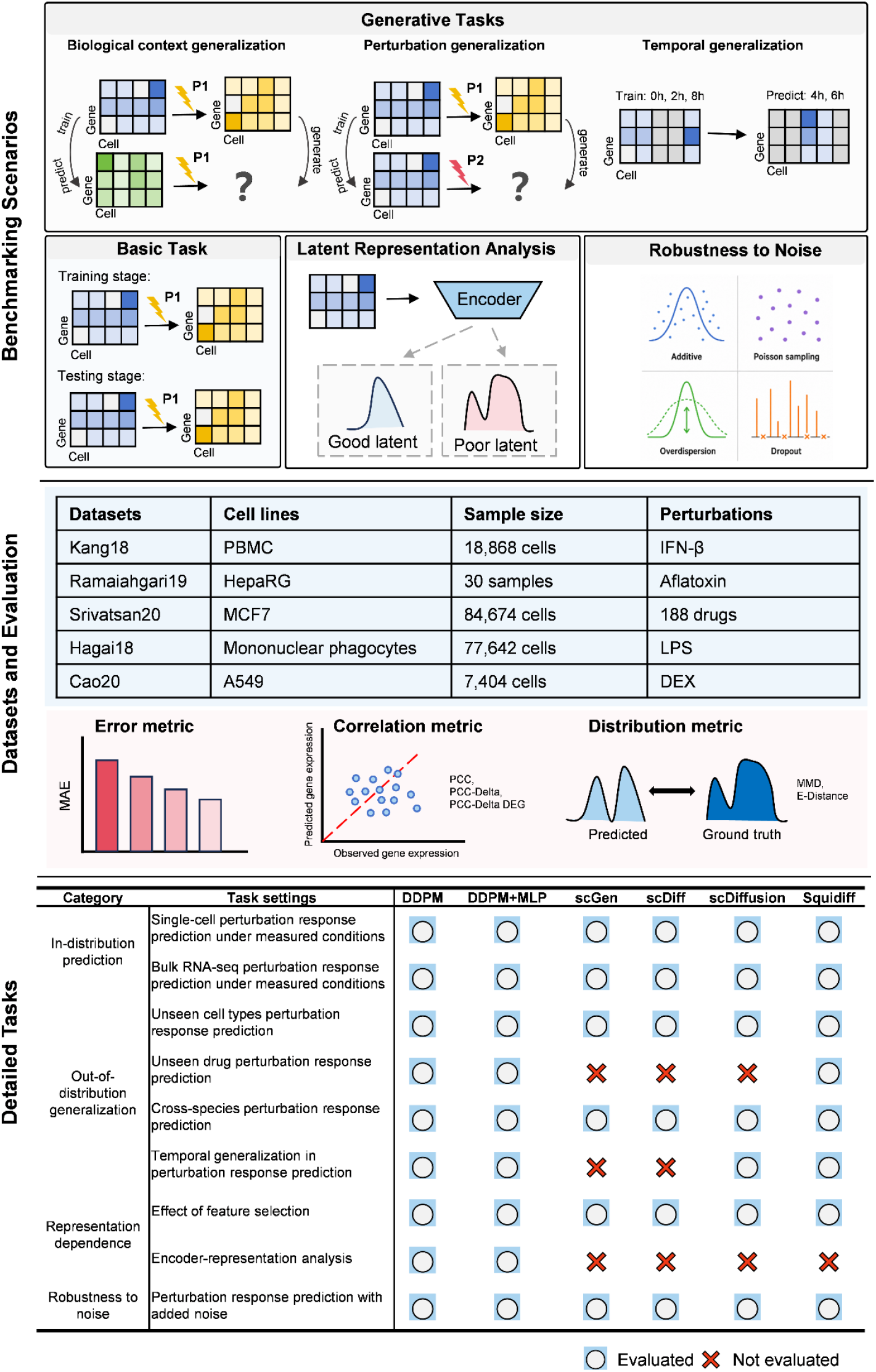
Framework of PertDiffBench. PertDiffBench evaluates diffusion-based models for perturbation response prediction (PRP) across in-distribution prediction, out-of-distribution generalization, representation-dependence analyses and noise-robustness tests. It includes several single-cell and bulk perturbation datasets. Model performance is assessed using error-based, correlation-based and distributional metrics, including MAE, Pearson correlation, PCC-Delta, DEG-based PCC-Delta, MMD and E-distance. The lower panel summarizes the benchmarked models, including DDPM, DDPM+MLP, scGen, scDiff, scDiffusion and Squidiff, and their applicable task settings; blue circles denote applicable settings and red crosses denote inapplicable settings.

PertDiffBench first evaluates prediction under measured perturbation conditions using two standard settings. The Kang18 dataset is used to test single-cell perturbation prediction in seen cell types, whereas the Ramaiahgari19 dataset is used to test bulk-level drug perturbation prediction. These tasks evaluate diffusion-based models against the VAE-based baseline scGen under standard perturbation response prediction settings, providing a reference point for the later analyses of generalization and robustness.

The benchmark then extends to unseen-condition prediction across three extrapolation settings. Biological-context generalization is evaluated through unseen cell-type prediction on Kang18 and cross-species prediction of LPS responses in the Hagai18 dataset. Perturbation generalization is evaluated on Srivatsan20 by testing prediction for unseen drug perturbations. Temporal generalization is evaluated on Cao20 through a temporal imputation task, which reflects the sparse time sampling common in cell differentiation and dynamic perturbation experiments and tests whether models can recover cellular states at unobserved intermediate time points. These settings assess not only the fit to observed perturbation patterns, but also the reliability of generalization across biological context, perturbation type and temporal dynamics.

Beyond task-level comparisons, PertDiffBench examines how data representation, model components and data noise affect prediction. We compare performance across feature dimensions and evaluate how different single-cell data representation methods work with the DDPM backbone, including STATE, scFoundation, scGPT, Tahoe-x1, Geneformer, Scimilarity, and scVI. These experiments ask whether performance is mainly determined by the diffusion backbone or depends strongly on data representation. Because single-cell data are sparse, overdispersed and affected by dropout and technical noise, PertDiffBench also includes a noise robustness analysis. We introduce Gaussian noise, Log-Normal noise, Poisson noise and Zero-Inflation to measure how performance changes under different data perturbations and to test whether the denoising premise of diffusion models leads to a practical robustness advantage.

At the model and metric levels, all models are evaluated with the same data processing pipeline, task splits and metrics. The primary metric is PCC-Delta, supplemented by PCC, MAE, MMD, E-distance and metrics based on top differentially expressed genes. Together, these metrics measure prediction quality at the levels of global expression state, perturbation response direction and key differentially expressed genes. PertDiffBench therefore addresses a diffusion model specific question: whether diffusion models are well suited to single-cell perturbation response prediction, and under which data conditions and task settings their strengths and limitations become apparent.

### Diffusion models do not consistently outperform scGen under seen perturbation conditions

To assess the baseline predictive performance of diffusion models under seen perturbation conditions, we compare diffusion-based models with the VAE-based model scGen in two standard perturbation response prediction tasks, one at the single-cell level and the other at the bulk level. In the Kang18 known-cell-type prediction task, scGen achieves the highest PCC-Delta and outperforms all diffusion-based models. Among the diffusion models, scDiffusion performs best, followed by Squidiff, DDPM, scDiff and DDPM+MLP, with progressively lower performance (Figure 2A, Supplementary Figure 1A and 1B). The bulk-level drug perturbation task shows a similar pattern. scGen and scDiffusion rank first and second, respectively, whereas the other diffusion models show weaker overall performance (Figure 2B, Supplementary Figure 1C). These results suggest that, under measured perturbation conditions, diffusion models do not consistently outperform classical VAE-based methods, and their performance varies substantially across model designs.

**Figure 2.**
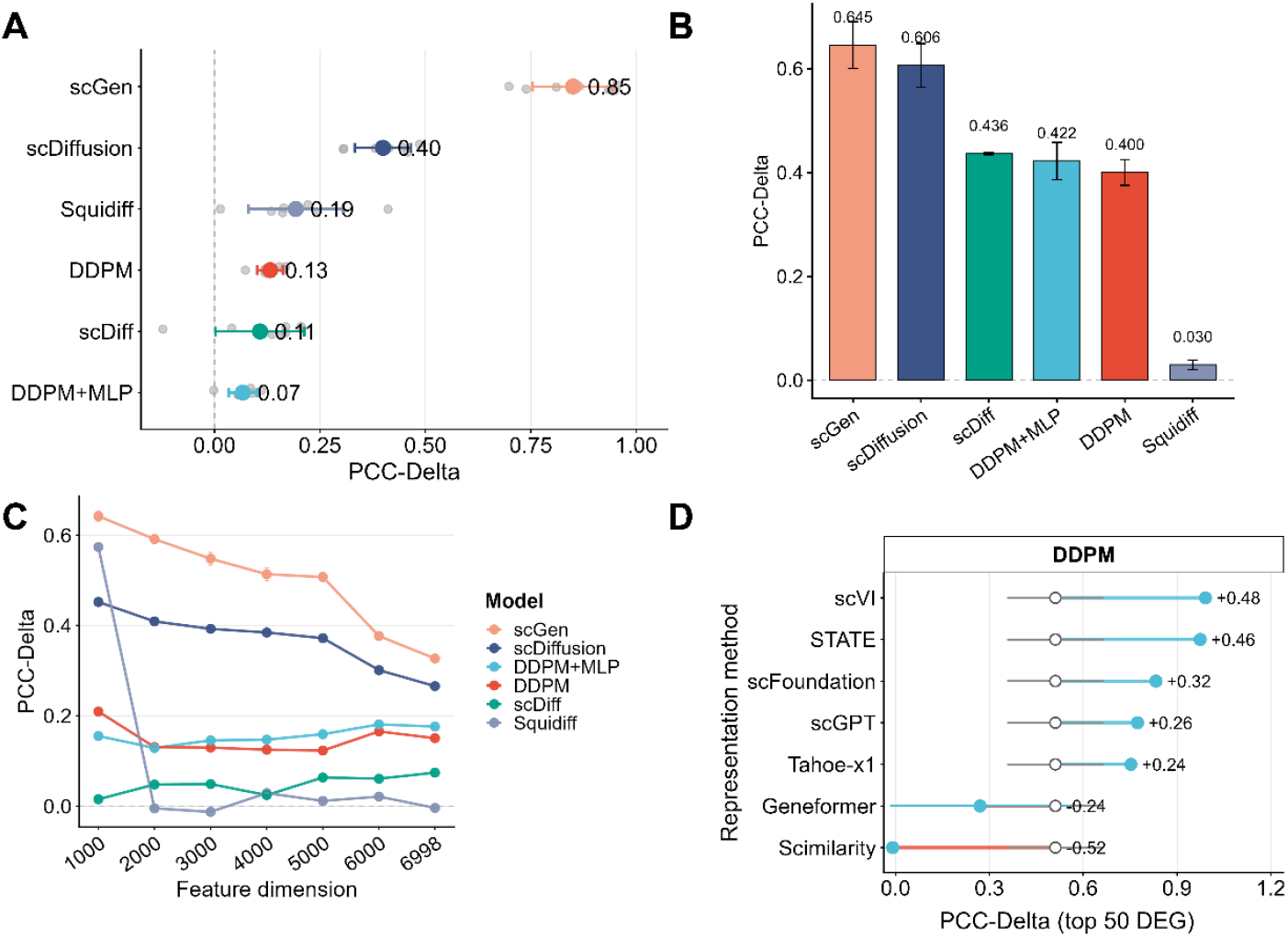
Baseline performance under seen perturbation conditions. (A) Single-cell perturbation response prediction under seen perturbation conditions. Model performance was evaluated across cell types using PCC-Delta. Gray points indicate performance for individual cell types, and colored points with error bars indicate the mean±s.d. across cell types. (B) Bulk-level drug perturbation response prediction under seen perturbation conditions. Models were compared in a drug-split task using PCC-Delta. (C) Effect of feature dimensionality on model performance in the CD4 T cell seen-condition prediction task. Models were evaluated using different numbers of input features. (D) Effect of encoder choice on DDPM performance in the CD4 T cell seen-condition prediction task.

We next examine how input feature dimensionality and representation affect predictive performance. As shown in Figure 2C, most models lose performance as the feature dimension increases. scGen remains the strongest model across all feature dimensions, followed by scDiffusion. DDPM, DDPM+MLP and scDiff stay at relatively low performance levels, whereas Squidiff performs well with low-dimensional features but declines rapidly as the feature dimension increases (Supplementary Figure 1D). For the known-cell-type prediction task, using CD4 T cells as the evaluation setting, we evaluated single-cell foundation models and classical representation models as encoders for DDPM. scVI and STATE produced the largest gains, followed by scFoundation, scGPT and Tahoe-x1, whereas Geneformer and Scimilarity reduced PCC-Delta on the top 50 differentially expressed genes and all genes (Figure 2D, Supplementary Figure 1E). The results suggest that the input representation may affect model performance more than the diffusion architecture itself.

Overall, diffusion models show measurable predictive performance under in-distribution perturbation conditions, but their performance is unstable and they do not generally outperform scGen. Performance depends strongly on feature dimensionality and data representation, indicating that representation learning is a key determinant of diffusion-based models for single-cell perturbation response prediction.

### Diffusion models show task-dependent generalization under unseen conditions

To assess model generalization under unseen conditions, we design extrapolation tasks that span cell type, species and perturbation type. In the unseen cell-type prediction task on the Kang18 dataset, we retain 25% of control cells from the held-out cell type as input and compare the predictive performance of different models. scGen achieves the best across PCC, PCC-Delta, MMD and MAE, followed by scDiffusion, whereas DDPM, DDPM+MLP, scDiff and Squidiff show weaker performance (Figure 3A). These results indicate that the VAE-based model scGen remains strong for unseen cell-type prediction, and that diffusion models do not consistently outperform scGen in this setting.

**Figure 3.**
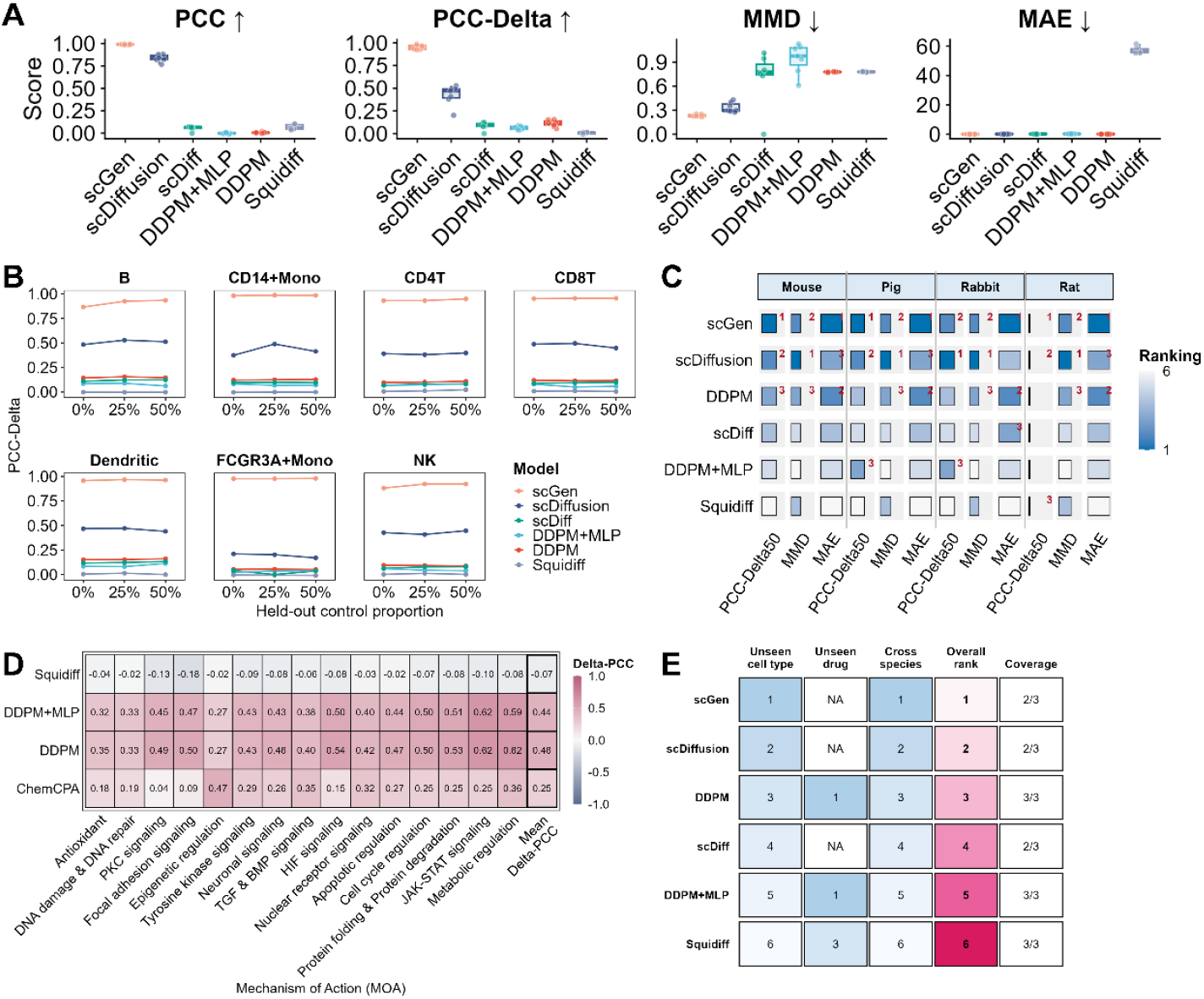
Model performance across out-of-distribution perturbation response prediction tasks. (A) Unseen cell-type prediction with 25% control cells from the held-out cell type provided as input. Models were evaluated using multiple metrics, including PCC, PCC-Delta, MMD and MAE. (B) Effect of control-cell proportion on unseen cell-type prediction. Models were evaluated with 0%, 25% and 50% control cells from the held-out cell type. PCC-Delta was compared across cell types and control-cell proportions. (C) Cross-species prediction of perturbation responses. Models were trained on a subset of species and tested on held-out species. Performance was evaluated using PCC-Delta, MMD and MAE. (D) Unseen mechanism-of-action prediction. (E) Summary of model rankings across unseen cell-type, unseen drug and cross-species prediction tasks. NA indicates models that were not evaluated for a given task because the model setting was not applicable. Coverage denotes the number of tasks included for each model.

We next examine how the proportion of control cells affects unseen cell-type prediction. As the fraction of control cells in the held-out cell type increases from 0% to 25% and 50%, PCC-Delta remains stable for most models (Figure 3B). scGen showed consistently high and stable performance across all held-out cell types. Among diffusion-based models, scDiffusion performed best overall, but its performance was lower for FCGR3A+ monocytes, indicating context-dependent generalization. Other diffusion-based models remained substantially weaker, with Squidiff performing close to zero across settings. These results suggest that control cells provide useful state references, but generalization is still shaped by the complexity of cell-type-specific responses and by model architecture.

We further evaluate cross-species prediction of LPS responses using the Hagai18 dataset. This task requires models to learn perturbation responses from a subset of species and predict cellular state changes in an unseen species, making it more challenging than cell-type transfer within the same species. scGen and scDiffusion show the strongest overall performance, DDPM performs at an intermediate level, and scDiff, DDPM+MLP and Squidiff perform less well (Figure 3C). These results show that cross-species generalization varies substantially across diffusion models, with only a few approaching the performance of the VAE-based baseline.

Finally, we evaluated cross-MOA unseen drug prediction, where models were trained on perturbations from other mechanisms of action and tested on held-out MOA groups. We compared DDPM, DDPM+MLP and Squidiff, and included ChemCPA, an adversarially regularized autoencoder-based model for compound-conditioned perturbation response prediction, as a task-specific baseline. DDPM and DDPM+MLP achieved the best overall PCC-Delta, with mean values of 0.46 and 0.44, respectively, outperforming ChemCPA across most MOA groups (Figure 3D). ChemCPA showed moderate performance but varied substantially across mechanisms, whereas Squidiff consistently produced negative PCC-Delta values. These results indicate that cross-MOA generalization remains highly model-dependent and mechanism-dependent, and that simple DDPM-based models can provide stronger performance than the compound-conditioned baseline in this setting.

Taken together, the unseen-condition tasks reveal the operating limits of different models. scGen performs consistently in unseen cell-type and cross-species prediction, suggesting that VAE-based latent perturbation modeling remains competitive in these extrapolation settings. The performance of diffusion-based models depends more strongly on the task and implementation. scDiffusion is relatively stable across several tasks, whereas DDPM baseline performs well in unseen MOA prediction. These results indicate that diffusion models do not provide a consistent advantage across unseen-condition settings (Figure 3E). Their generalization is shaped by the biological context, the extrapolation setting, and the input embedding.

### Diffusion models show model-specific behavior in temporal imputation

To assess whether diffusion models can infer missing time points, we design a temporal imputation task using single-cell omics data from DEX-treated A549 cells collected at 0, 2, 4, 6, 8, and 10 h. The models are trained on the remaining time points and used to predict the held-out 4 h and 6 h cell states (Figure 4A and B). The models show clear differences in this setting (Figure 4C). DDPM shows the strongest overall performance at both held-out time points. It achieves the lowest MMD and MAE and the highest PCC-Delta, indicating better recovery of both the expression distribution and perturbation-induced changes at 4 h and 6 h. scDiffusion achieves the highest PCC, suggesting accurate reconstruction of overall expression profiles, but it is weaker than DDPM on PCC-Delta and distribution-level metrics. DDPM+MLP and Squidiff perform poorly overall, with near-zero PCC and higher MMD and MAE. These results suggest that, for temporal imputation, a simple DDPM backbone can outperform more complex diffusion variants.

**Figure 4.**
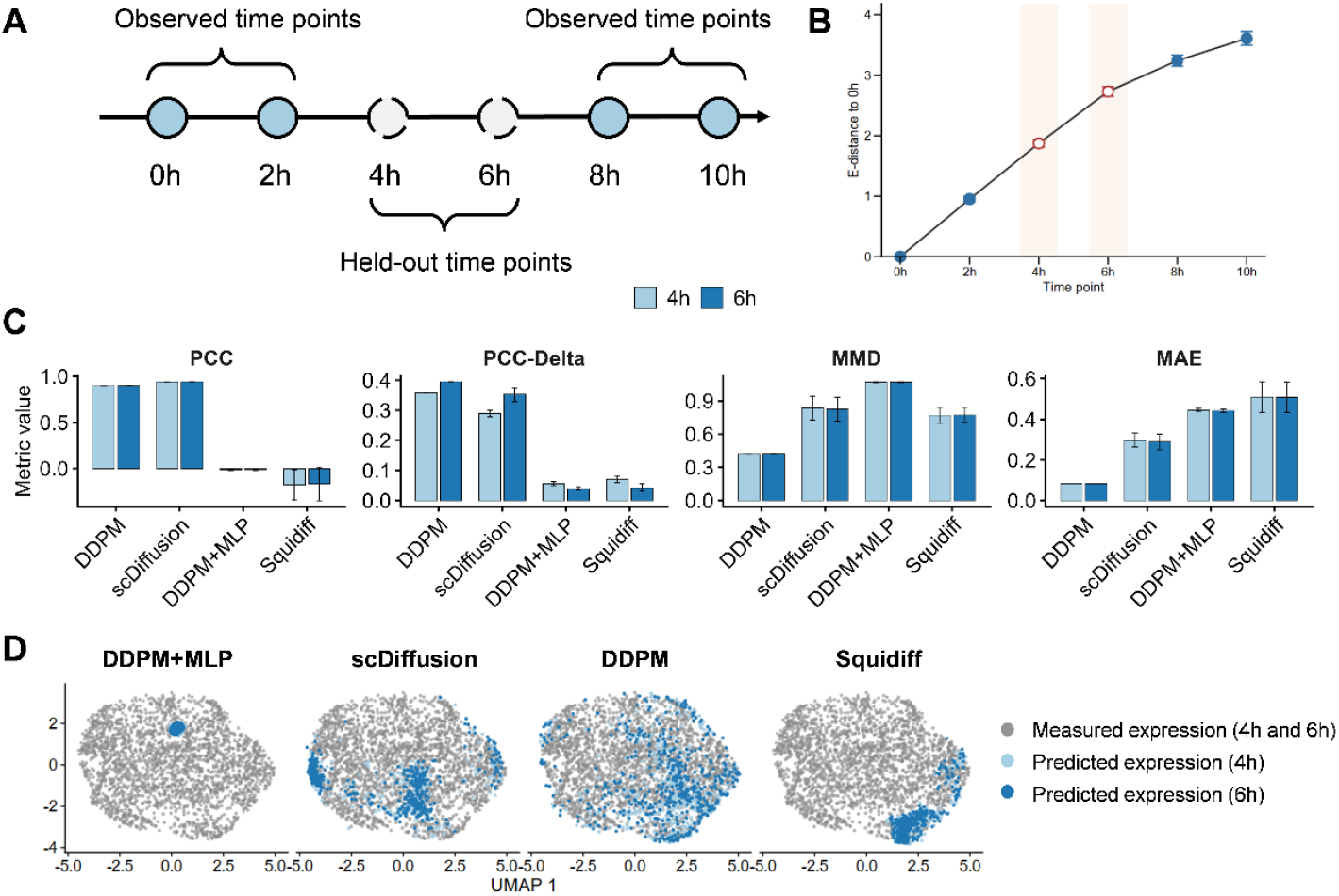
Temporal interpolation of perturbation responses at held-out time points. (A) Schematic of the temporal interpolation task. (B) E-distance from each time point to the 0 h baseline, showing the progression of transcriptional changes over time. Shaded regions indicate the held-out time points used for model evaluation. (C) Temporal imputation performance on DEX-treated A549 single-cell omics data. Models were trained using the observed time points and evaluated on the held-out 4 h and 6 h cell states. Performance was measured using PCC, PCC-Delta, MMD and MAE. (D) UMAP visualization of predicted cells for the held-out 4 h and 6 h time points across three repeated runs. Gray points represent reference cells, and blue points represent model-generated cells, with light blue indicating 4 h and dark blue indicating 6 h.

The UMAP visualization reveals distinct patterns for each model (Figure 4D). DDPM produces cells spread across a wider region, but local neighborhoods are less coherent. Cells generated by scDiffusion closely follow the reference distribution. Squidiff predictions are concentrated in specific branches, while DDPM+MLP forms a tight cluster of predicted cells, consistent with strong mode collapse.

These results show that diffusion models can impute unobserved intermediate time points, but their ability to recover temporal structure varies across implementations. DDPM better reconstructs the overall expression distribution.

### Failure modes and sensitivity analysis of diffusion-based perturbation response prediction

We next asked which factors most strongly affected diffusion-based perturbation response prediction. We first examined the role of input representation by pairing DDPM with a set of pretrained and classical single-cell encoders in the unseen cell-type task. Under the 25% control-cell setting, DDPM performance varied markedly across encoders and target cell types (Figure 5A). scVI, a classical VAE-based representation model, achieved the strongest overall performance, followed by STATE. By contrast, larger pretrained transcriptomic language models, including scGPT, scFoundation, and Tahoe-x1 showed intermediate performance and rarely outperformed scVI. Scimilarity and Geneformer performed poorly across most cell types, with PCC-Delta values close to zero in several cases. Performance also differed across target cell types, with B cells, CD8 T cells, dendritic cells and NK cells being more predictable than FCGR3A+ monocytes. These results suggest that effective perturbation response prediction does not simply follow model scale. Instead, a compact VAE-based representation such as scVI can provide a more suitable latent space for DDPM-based generation than several larger pretrained encoders.

**Figure 5.**
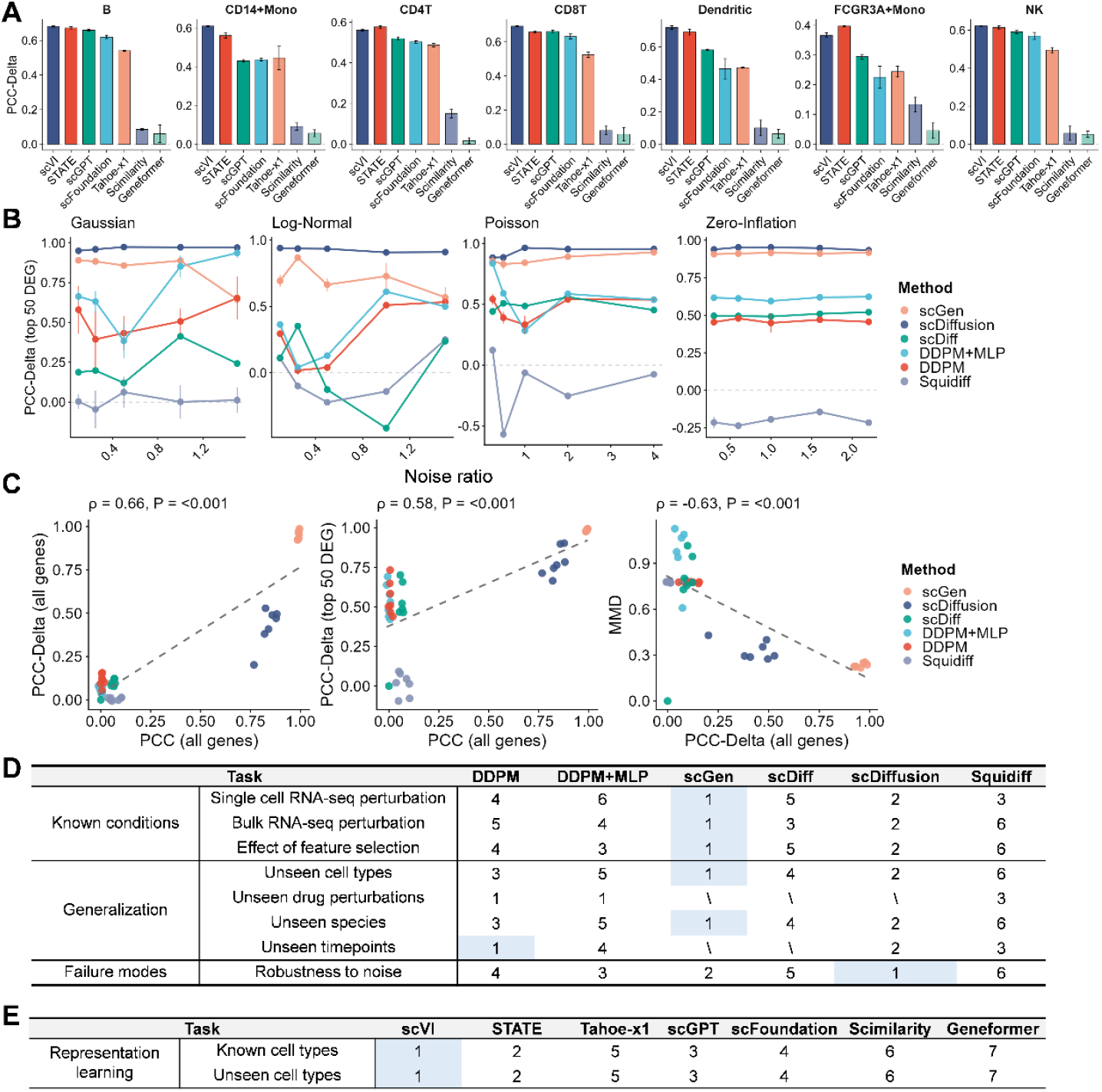
Sensitivity analysis. (A) Effect of input representation on DDPM-based prediction across held-out cell types. DDPM was paired with pretrained and classical single-cell encoders, and performance was assessed using PCC-Delta. (B) Robustness of perturbation response prediction models to input noise. Gaussian, Log-Normal, Poisson and Zero-Inflation noise were added at increasing noise ratios, and performance was measured using PCC-Delta on the top 50 differentially expressed genes. (C) Agreement between evaluation metrics across models and tasks. Scatter plots compare PCC with PCC-Delta, PCC with PCC-Delta on the top 50 differentially expressed genes, and PCC-Delta with MMD. Dashed lines show fitted linear trends, and ρ and p values indicate Spearman correlation statistics. (D) Overall ranking of model performance across benchmark tasks, including known-condition prediction, generalization and robustness to noise. Lower ranks indicate better performance, and blue shading marks the best-performing model for each task. (E) Ranking of representation models used as DDPM encoders for known and unseen cell-type prediction. Lower ranks indicate better performance, and blue shading marks the top-ranked encoder.

We next examined whether these encoder-dependent differences were associated with perturbation-aware structure in the embedding space. To account for differences in prediction difficulty across held-out cell types, we centered both representation metrics and PCC-Delta values within each held-out cell type. In this analysis, condition kNN purity showed a positive association with PCC-Delta on the top 50 differentially expressed genes, whereas the association was weaker for PCC-Delta across all genes (Supplementary Figure 3). By contrast, perturbation signal ratio showed little association with either metric. These results suggest that DDPM performance in the unseen-cell-type setting is more closely related to the preservation of local Control/IFN-perturbed neighborhood structure than to the magnitude of the global IFN perturbed–Control centroid shift alone.

We then evaluated model robustness by adding Gaussian, Log-Normal, Poisson and Zero-Inflation noise to the input expression profiles. scDiffusion was the most stable model across noise types and noise ratios, maintaining high PCC-Delta on the top 50 differentially expressed genes (Figure 5B). scGen also remained competitive, although its performance varied more under Gaussian and Log-Normal noise. DDPM and DDPM+MLP showed stronger dependence on the noise distribution. DDPM+MLP performed well under high Gaussian noise but was less stable under Log-Normal and Poisson noise, whereas DDPM showed moderate performance across most settings. scDiff was sensitive to the type of noise, with a marked drop under log-normal noise. Squidiff was the least robust model overall, producing near-zero or negative PCC-Delta under several noise settings.

We also tested whether model performance was sensitive to the ordering of input genes. We compared a cluster-based gene ordering with a random gene shuffle, with the same order applied consistently to training and validation data. Most models showed limited sensitivity to gene order, including scGen, DDPM+MLP and Squidiff (Supplementary Figure 4). By contrast, DDPM and scDiffusion showed significant differences between the two orderings, although the direction of the effect was model dependent. These results indicate that gene order is not a universal driver of performance, but can affect models whose implementation implicitly or explicitly uses feature organization.

Across the benchmark, commonly used evaluation metrics were correlated but not interchangeable (Figure 5C). PCC was positively correlated with both all-gene PCC-Delta and top-50-DEG PCC-Delta, whereas MMD was negatively correlated with PCC-Delta, consistent with their opposite performance directions. The overall ranking further showed that no single model dominated every task (Figure 5D). scGen and scDiffusion ranked highest in several known-condition and generalization settings, while DDPM-based models remained competitive in selected tasks, including unseen drug perturbation response prediction. The encoder comparison showed a similar pattern at the representation level, with scVI and STATE ranking above other pretrained models in both known and unseen cell-type settings (Figure 5E). Together, these analyses indicate that model performance depends not only on the generative backbone, but also on the input representation, noise structure and evaluation metric.

## Discussion

Our benchmark shows that diffusion models are not a default improvement over established generative baselines for single-cell perturbation response prediction. Across the tasks examined here, scGen remained the strongest model in the seen-condition setting, while scDiffusion was the most competitive diffusion-based method. Other diffusion variants showed more variable performance. This result does not rule out the value of diffusion models, but it argues against treating the diffusion backbone itself as sufficient for improved prediction.

A recurring pattern across the benchmark is the importance of representation. In the feature-dimensionality analysis, most models lost accuracy as the input space became larger. scGen remained robust across feature dimensions, followed by scDiffusion, whereas DDPM, DDPM+MLP and scDiff stayed at relatively low performance levels. Squidiff performed well with low-dimensional features but declined rapidly when more genes were included. These results point to a practical issue for diffusion-based modelling in omics: denoising in gene-expression space may be sensitive to the scale and structure of the input features. Unlike images, gene-expression matrices do not have an obvious spatial neighbourhood structure, and dependencies between genes are often nonlocal and context dependent. The effectiveness of a diffusion model may therefore depend heavily on whether the input representation exposes a learnable structure for the denoising process.

The encoder experiments support this view. Pretrained representations did not provide a uniform benefit. In the unseen cell-type task, scVI and STATE gave stronger and more stable results than several larger trained encoders. This suggests that encoder quality should not be judged only by pretraining scale. For perturbation response prediction, a useful representation needs to preserve cell-state variation and perturbation-relevant expression structure in a form that is compatible with the downstream generative model.

Generalization performance also depended on the biological setting. In unseen cell-type prediction, scGen remained the strongest overall model, followed by diffusion-based methods. Adding control cells from the held-out cell type produced only modest changes, and the effect varied across cell types. This implies that cell-type extrapolation is not determined only by the amount of control information available, but also by how transferable the target cell state is from the training conditions. In cross-species prediction, scGen and scDiffusion alternated as the best-performing methods, with DDPM usually ranking below them. Thus, diffusion-based models can be competitive in some out-of-distribution settings, but the advantage is not consistent across tasks.

Temporal imputation showed a different behavior. In this task, the DDPM baseline applied directly to the original gene-expression space outperformed DDPM+MLP and scDiffusion, whereas Squidiff showed the weakest performance. This result suggests that temporal imputation is particularly sensitive to how intermediate cellular states are represented. Direct modeling in gene-expression space may preserve subtle, time-dependent changes in gene abundance, while DDPM+MLP compresses the input through a three-layer MLP into a 256-dimensional latent space, which may remove part of this information. These results indicate that the relative performance of diffusion models depends not only on the diffusion backbone, but also on whether the chosen representation preserves the structure required by the task.

The robustness analysis further shows that model performance depends on the type of noise introduced into the input data. Log-Normal overdispersion produced the largest decrease in top-DE-gene PCC-Delta, whereas dropout had a more limited effect. This difference may reflect the fact that overdispersion alters gene-level variability and response magnitude, both of which are central to perturbation response prediction, while dropout mainly increases sparsity. Robustness to one form of noise therefore does not necessarily imply robustness to another. Perturbation-model benchmarks should include noise settings that capture distinct sources of technical and biological variation, rather than relying on a single generic corruption test.

Taken together, these results suggest that the main determinant of performance is not the diffusion backbone alone, but its interaction with representation, feature space and task design. Diffusion models may be useful for single-cell perturbation response prediction, but their current advantage is conditional rather than universal. Future methods may need more omics-specific inductive biases, better task-aligned representations, and explicit use of regulatory and pathway structure to make the denoising process better matched to gene-expression data.

This study has several limitations. The benchmark focuses mainly on transcriptomic perturbation responses and does not yet cover the full range of multimodal perturbation readouts. The evaluated methods represent current diffusion-based approaches, but they do not exhaust all possible architectures. Some failure-mode analyses, including sensitivity to feature ordering, require further investigation. Finally, quantitative benchmarks can compare model behavior across controlled settings, but they cannot replace biological validation. Future work should extend these evaluations to larger perturbation atlases, multimodal measurements and more stringent out-of-distribution settings, with particular attention to separating the effects of representation, model architecture and task design.

## Datasets and Code Availability

https://github.com/ZijunSong/PertDiffBench

## Author Contributions

L.X. and Y.X. conceived the study and designed the PertDiffBench framework. J.Z.S., Y.X., Z.S., and W.J. developed the benchmark tasks, evaluation settings and analysis workflow. J.Z.S. implemented the benchmarking pipeline and performed model training and evaluation. Y.X. and J.L. performed data analysis. Y.X., J.Z.S., L.X., Z.S., W.J., J.L., and C.S. wrote the manuscript. All authors reviewed and approved the final manuscript.

## Funding

This work was supported by National Key R&D Program of China, 2025YFA1309400.

## Competing interests

The authors declare no competing interests.

## Methods

### Datasets and preprocessing

We used Kang18 and Ramaiahgari19 to benchmark response prediction in known perturbation contexts, covering single-cell RNA-seq and bulk RNA-seq data, respectively (Kang, et al., 2018; Ramaiahgari, et al., 2019). For unseen-context prediction, we used Srivatsan20 for compound perturbation generalization, Kang18 for cross-cell-type prediction and Hagai18 for cross-species transfer (Hagai, et al., 2018; Srivatsan, et al., 2020). A549 time-resolved datasets were used to evaluate temporal imputation (Cao, et al., 2020). For all datasets, expression matrices were centered and log-normalized before model training and evaluation. The number of genes used for each benchmark setting is provided in Supplementary Table 1. Task-specific transformations and train-test split procedures are described in the corresponding Methods sections.

### Benchmarking tasks and experiment settings

To systematically evaluate perturbation-response prediction, we organized the benchmark into four scenarios: known-condition prediction, unseen-condition generalization, robustness analyses, and representation-learning benchmarking. These settings were designed to cover both standard in-distribution prediction and biologically meaningful distribution shifts, including transfer across perturbations, cellular contexts, species, and treatment times.

#### Known-condition prediction

Known-condition prediction was used to assess whether models could generate post-perturbation expression profiles for perturbation contexts observed during training. For the scRNA-seq benchmark, we used seven cell types from the Kang PBMC dataset: B cells, CD14+ monocytes, CD4 T cells, CD8 T cells, dendritic cells, FCGR3A+ monocytes and natural killer cells. Each cell type was analysed separately, and models were evaluated on held-out control and perturbed profiles from the same cell type and perturbation context.

For the bulk RNA-seq benchmark, we used the Ramaiahgari19 dataset from LINCS, which contains expression profiles before and after aflatoxin treatment across ten dose levels. The dataset included 12 control samples and 30 treated samples, with three replicates for each dose. We generated three random train-test splits. In each split, six control samples and 20 treated samples, two replicates per dose, were used for training, while the remaining six control samples and ten treated samples, one replicate per dose, were held out for testing. Models were evaluated by their ability to predict the held-out treated expression profiles from control inputs under perturbation conditions represented during training.

#### Biological-context generalization

For cross-cell-type prediction, each cell type was held out in turn, and the remaining cell types were used for training. For each held-out cell type, we used two protocols. In the fully out-of-distribution setting, all samples from the held-out cell type were excluded from training. In the partially shared setting, control cells from the held-out cell type were included during training, but the corresponding perturbed cells were kept for testing. This design allowed us to distinguish transfer to an entirely unseen cellular context from prediction in a target context where baseline expression profiles were available.

The cross-species task followed the same leave-one-context-out design. Each species was held out in turn, with the remaining species used for training under the same perturbation setting. Models were then evaluated on the held-out species, allowing us to test whether perturbation responses learned in one set of species could be transferred to another.

#### Perturbation generalization

Perturbation generalization was evaluated in two settings: intra-MOA unseen drug prediction and cross-MOA unseen mechanism prediction. For intra-MOA prediction, each mechanism-of-action (MOA) category was analysed independently. Within each MOA, drugs were split into training and test sets. Models were trained to predict drug-treated expression profiles from matched control profiles using the training drugs, and were evaluated on held-out drugs from the same MOA. This analysis was performed across 15 MOA-specific splits.

For cross-MOA prediction, we used a leave-one-MOA-out design. In each split, drugs from one MOA were held out for testing, and drugs from the remaining MOAs were used for training. This procedure was repeated for all 15 MOA categories. Because this task requires prediction for drugs from unseen mechanisms, we restricted the benchmark to models that support drug-conditioned generation. Drug information was provided according to the input format required by each model.

For DDPM, DDPM+MLP and Squidiff, compound identity and dose were encoded jointly as a structure–dose representation. Specifically, each compound was represented by a 1,024-dimensional RDKit Morgan feature fingerprint using a radius of 2; for multi-component SMILES strings, component-level fingerprints were averaged. The resulting molecular fingerprint was then scaled by log_10_(dose + 1), so that dose information modulated the structural representation rather than being provided as an independent categorical variable. During testing, held-out compounds were encoded from their SMILES strings using the same procedure, with the corresponding dose summarized by the median observed dose for that condition. Squidiff used the same structure–dose representation in the cross-MOA experiments. ChemCPA used a different encoding scheme. Each canonical SMILES string was converted into a 2,048-bit RDKit Morgan fingerprint with radius 2, while the dose was encoded separately by ChemCPA’s learned dose network before decoding.

#### Time-point generalization

For temporal imputation, we used a time-resolved single-cell RNA-seq dataset of A549 cells treated with dexamethasone (DEX), with cells collected at 0 h, 2 h, 4 h, 6 h, 8 h and 10 h after treatment. Models were trained on observed response states at 0 h, 2 h, 8 h and 10 h, and evaluated on the held-out intermediate time points at 4 h and 6 h. This task assesses whether a model can recover unseen intermediate transcriptional states along a perturbation-induced temporal trajectory.

The models used different latent representations for intermediate-state generation. The direct-expression DDPM baseline was trained directly in gene-expression space, and the held-out 4 and 6 h states were estimated by linear interpolation between the observed 2 and 8 h expression profiles, without an auxiliary encoder or decoder. DDPM+MLP used the same interpolation scheme in a learned 256-dimensional latent space before decoding the interpolated states back to gene-expression space.

For Squidiff, temporal imputation was performed in its semantic latent space. In the default addition mode, 2 h cells were encoded into 60-dimensional semantic representations, and a global transition vector was estimated as the difference between the mean 8 h and mean 2 h latent states. Intermediate states were generated by moving the 2 h latent representation along this transition direction according to the target time point, followed by DDIM decoding. Cell-level stochastic variation was retained during decoding. We also evaluated a linear-interpolation variant of Squidiff, in which 2 and 8 h semantic latents were mixed directly on a per-cell basis before decoding.

For scDiffusion, temporal imputation was performed in a 128-dimensional VAE latent space using classifier-guided gradient interpolation rather than direct latent mixing. The model was trained as a three-component pipeline on the observed time points at 0 h, 2 h, 8 h and 10 h. First, a VAE initialized from a pretrained annotation encoder–decoder was used to map expression profiles into latent representations. A diffusion model was then trained on these latent codes, and a time-point classifier was trained to distinguish the four observed treatment times from VAE-encoded cells. To generate the held-out 4 h and 6 h states, cells from the 2 h state were encoded into the VAE latent space and diffused to an intermediate noise level before reverse sampling. During reverse diffusion, classifier guidance was applied jointly with respect to the 2 h and 8 h time labels, thereby steering samples between the early and late observed response states. The relative guidance strengths were controlled by predefined interpolation weights assigned to the two endpoint labels. For the 4 h prediction, balanced guidance toward 2 h and 8 h was used, whereas the 6 h prediction used stronger guidance toward the 8 h state. Classifier guidance was applied during the final portion of the reverse-diffusion trajectory, and the resulting latent samples were decoded back into gene-expression space.

Unlike the linear-interpolation baselines, scDiffusion used observed time labels explicitly through the auxiliary classifier during both training and sampling. The temporal-imputation experiments did not use SMILES or dose embeddings for Squidiff, and the DDPM-family models were not provided with explicit time-point embeddings. Thus, for these baselines, temporal information was inferred from the latent geometry between observed early and late response states.

### Baseline models

The benchmark included two implemented DDPM baselines, DDPM and DDPM+MLP, together with published models, including scGen, scDiff, scDiffusion and Squidiff.

#### DDPM

The direct-expression DDPM baseline used a conditional denoising diffusion model operating on preprocessed gene-expression vectors. Let *x*_0_ ∈ ℝ^*G*^ denote a perturbed expression profile and *c* ∈ ℝ^*G*^ denote the corresponding control-profile input. The forward diffusion process was defined as

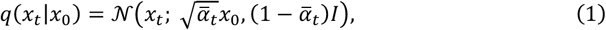

Where 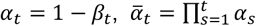, and 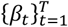 followed a linear variance schedule.

The denoising model *𝜖*_*θ*_(*x*_*t*_, *c, t*) was parameterized by a compact conditional multilayer perceptron. The noisy target profile and the control profile were concatenated and projected into a hidden representation, while the diffusion timestep was encoded with a sinusoidal timestep embedding. These two representations were combined before the final noise-prediction layers:

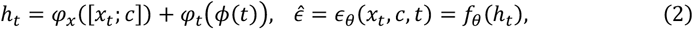

where [*x*_*t*_; *c*] denotes concatenation along the gene dimension, *ϕ*(*t*) is the sinusoidal timestep embedding, and *φ*_*x*_, *φ*_*t*_ and *f*_*θ*_ are fully connected transformations. This architecture provides conditional generation from control to perturbed expression space without introducing convolutional assumptions over the gene order.

The model was trained with the standard DDPM noise-prediction objective,

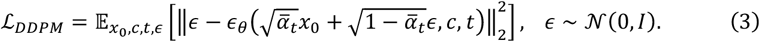

At inference, samples were generated by conditional reverse diffusion initialized from Gaussian noise and conditioned on the control profile *c*. Generated profiles were evaluated directly in the same preprocessed expression space used for training. The default configuration used *T* = 1,000 diffusion steps, a linear schedule from *β*_1_ = 10^−4^ to *β*_*T*_ = 0.02, and a hidden width of 1,024. Optimization used AdamW with cosine learning-rate annealing and gradient clipping.

#### DDPM+MLP

The DDPM+MLP baseline extended the direct-expression DDPM by moving the diffusion process from gene-expression space to a learned latent space. Let *E*_*ϕ*_ and *D*_*ω*_ denote an MLP encoder and decoder, respectively. For a control profile *c* ∈ ℝ^*G*^ and a perturbed target profile *x*_0_ ∈ ℝ^*G*^, the corresponding latent representations were

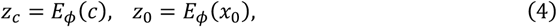

where *z*_*c*_, *z*_0_ ∈ ℝ^*L*^. Diffusion was applied to the perturbed latent representation *z*_0_, while the control latent representation *z*_*c*_ was used as the conditioning input.

The latent denoising network followed the same conditional noise-prediction principle as the direct-expression DDPM, but operated on *L*-dimensional latent variables rather than *G*-dimensional expression profiles:

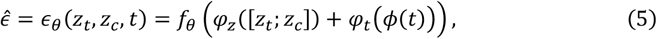

where *z*_*t*_ is the noisy perturbed latent variable, [*z*_*t*_; *z*_*c*_] denotes latent concatenation, *ϕ*(*t*) is the sinusoidal timestep embedding, and *φ*_*z*_, *φ*_*t*_ and *f*_*θ*_ are fully connected transformations. This design tests whether a compact latent representation improves conditional denoising by reducing the dimensionality of the generation space.

Training used the latent-space analogue of the DDPM noise-prediction objective:

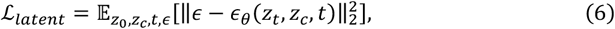

with *z*_*t*_ obtained by applying the forward diffusion process to *z*_0_. At inference, a perturbed latent profile was sampled by conditional reverse diffusion from Gaussian noise given *z*_*c*_, and the generated latent representation was decoded back to expression space:

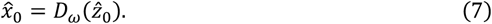

The generated expression profiles were evaluated in the same preprocessed expression space as the other baselines.

In the main benchmark, direct-expression DDPM and DDPM+MLP were evaluated as separate baselines to distinguish diffusion in expression space from diffusion in a learned latent space. Encoder-based representation experiments used a separate latent-diffusion setting in which pretrained or separately learned cellular embeddings were supplied to the downstream diffusion module; these experiments were therefore analysed as representation-learning benchmarks rather than as the main DDPM+MLP baseline.

#### Published baselines

scGen was used as the VAE-based perturbation response prediction baseline. It learns a latent representation of single-cell expression and estimates perturbation responses as shifts from control to perturbed states in the latent space. The diffusion-based published baselines, including scDiff, scDiffusion and Squidiff, were evaluated using the same task definitions, train-test splits and processed expression matrices as the implemented DDPM baselines whenever their model interfaces supported the required conditioning variables. For drug-response tasks, molecular structure and dose information were provided only to models whose native architecture supported these conditioning inputs.

Each published baseline was run with the configuration required by its original implementation. scGen was trained for up to 100 epochs with batch size 32 and early stopping with a patience of 25 epochs. scDiff was trained with the benchmark configuration for up to 1,000 epochs using a fixed data-split seed. scDiffusion was trained as a three-component pipeline consisting of a variational autoencoder, a diffusion model and a classifier; the VAE, diffusion and classifier components were each trained according to the corresponding benchmark configuration, with 10,000 optimization steps or iterations for the main components. In the repeated-generation setting, the trained scDiffusion components were reused and stochastic generation was repeated three times for evaluation. Squidiff was trained with its benchmark configuration using a learning rate of 10^−4^, 1,000 diffusion steps and batch size 128, without early stopping. This design kept data preprocessing, task splits and evaluation metrics unified across models, while preserving the model-specific optimization procedures needed for each published method.

### Representation-learning benchmark

For the representation-learning benchmark, cellular embeddings were first extracted with each encoder and then kept fixed during downstream perturbation response prediction. The downstream predictor was held constant across encoders and consisted of a latent conditional DDPM followed by an MLP decoder. The latent DDPM generated perturbed-state embeddings from control-state embeddings, and the decoder mapped the generated embeddings back to gene-expression space before metric computation. The model was trained with a combined objective that included the latent diffusion loss and a decoder reconstruction loss.

The evaluated encoders included scVI, STATE, Tahoe-x1, scGPT, scFoundation, Geneformer and Scimilarity. For each encoder, input preprocessing and gene handling followed the requirements of the corresponding representation method, including model-specific normalization, gene-vocabulary alignment or tokenization where applicable. Encoder dimensionalities and checkpoint choices followed the released benchmark configuration. This design allowed the benchmark to compare the practical utility of different cellular representation spaces while keeping the downstream generative model, task splits and evaluation metrics fixed.

In the main baseline comparison, DDPM and DDPM+MLP refer to the direct-expression diffusion baseline and the autoencoder-based latent diffusion baseline, respectively. In the encoder benchmark, by contrast, only the upstream cellular embedding was varied, whereas the downstream latent DDPM-decoder module was fixed across encoders.

### Perturbation-aware embedding structure analysis

To examine whether encoder-dependent DDPM performance was associated with perturbation-relevant structure in the embedding space, we quantified two perturbation-aware representation metrics for each encoder and held-out cell type. For each encoder, embeddings from all held-out cell types were pooled and standardized before PCA. We retained up to 50 principal components; for lower-dimensional embeddings such as scVI, all available dimensions were used. Metrics were then computed within each encoder-specific PCA space.

Condition k-nearest-neighbor (kNN) purity was used to quantify local perturbation-state structure. For each cell, we identified its 15 nearest neighbors in the embedding space and calculated the fraction of neighbors with the same perturbation condition, Control or IFN. The mean value across cells was used as the condition kNN purity for that encoder–held-out-cell-type pair.

Perturbation signal ratio was used to quantify global IFN perturbed–Control separation relative to within-condition variation. For each held-out cell type, we computed the centroids of Control and IFN perturbed cells in the embedding space. The perturbation signal ratio was defined as the Euclidean distance between the IFN perturbed and Control centroids divided by the square root of the mean within-condition variance, where within-condition variance was calculated from the squared distances of cells to their corresponding condition centroid.

To account for differences in prediction difficulty across held-out cell types, both representation metrics and DDPM performance values were centered within each held-out cell type before correlation analysis. DDPM performance was measured using PCC-Delta on the top 50 differentially expressed genes and, separately, on all genes. Associations between centered representation metrics and centered DDPM performance were evaluated using Spearman correlation across all encoder-held-out-cell-type pairs. Reported P values are nominal and were not adjusted for multiple testing.

### Robustness analyses

For the feature-dimensionality analysis, genes were ranked once by highly variable gene selection after library-size normalization and log1p transformation. Nested gene panels were then constructed by retaining the top-ranked genes at increasing feature dimensions (1,000, 2,000, 3,000, 4,000, 5,000 and 6,000 genes). When the aligned expression space allowed, an additional 6,998-gene panel was included. All panels were generated from the same underlying expression matrix so that cell assignments, perturbation labels and train-test splits were identical across feature dimensions.

For the noise-robustness analysis, we focused on CD4 T cells, the largest subpopulation in the Kang PBMC dataset. Noise was added separately to the training and validation data. For each noise type and severity level, models were trained from scratch and tested under the same conditional generation setting used for the clean data.

We assessed four types of input noise. Additive Gaussian noise was applied by adding independent zero-mean Gaussian noise to each expression value and truncating negative values to zero, with noise standard deviation *σ* ∈ {0.1, 0.25, 0.5, 1.0, 1.5}. Log-normal overdispersion was used to mimic excess cell-to-cell variability: gene-specific mean expression was estimated from the data, latent expression rates were sampled from a log-normal distribution with coefficient of variation *c*, and observed values were sampled from a Poisson distribution around these rates, with *c* ∈ {0.1, 0.25, 0.5, 1.0, 1.5}. Poisson noise was used to model depth-dependent sampling noise by drawing corrupted entries from a Poisson distribution with rate proportional to the original expression value and a sequencing-depth factor *d* ∈ {0.25, 0.5, 1.0, 2.0, 4.0}. Zero inflation was used to mimic dropout by assigning gene-specific dropout probabilities as a decreasing logistic function of gene abundance, with low-abundance genes assigned higher dropout probabilities than high-abundance genes. A global severity factor *s* ∈ {0.3, 0.6, 1.0, 1.6, 2.2} scaled these dropout probabilities, and entries were masked by independent Bernoulli draws.

### Feature ordering analysis

For the random-shuffle setting, gene columns were randomly permuted. For the cluster-reorder setting, genes were ordered by average-linkage hierarchical clustering of the gene-gene Pearson correlation matrix computed from log-normalized training data, with distance defined as 1 minus the correlation coefficient. In both settings, only the column order was changed; expression values were left unchanged, genes were not aggregated, and the same order was used for training and validation data.

### Evaluation metrics

Model outputs were evaluated at the distribution level. For a condition *c*, let 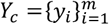 denote the held-out real perturbed profiles, 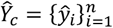 the generated perturbed profiles, and *B*_*c*_ the corresponding control or baseline profiles. Their mean profiles are denoted by 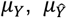 and *μ*_*B*_, respectively. Unless otherwise specified, all vector operations were performed across genes.

The primary response-level metric was PCC-Delta, which measures agreement between the predicted and observed perturbation effects. The observed and predicted effects were defined as

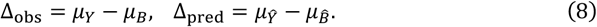

PCC-Delta was then computed as the Pearson correlation

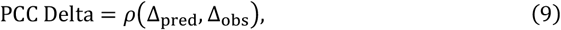

where, for two gene-wise vectors a and b,

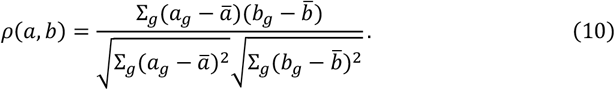

PCC-Delta therefore evaluates whether a model recovers the direction and relative magnitude of the perturbation-induced transcriptional response, rather than only reconstructing global expression levels.

Pearson Delta on differentially expressed genes was computed by restricting Delta PCC to the top *k* genes ranked by |Δ_obs_|:

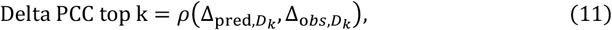

where *D*_*k*_ denotes the top k real response genes. We reported *k* = 20, 50 and 100, with *k* = 50 used as the primary top-DE subset when a single top-DE result was shown.

Mean-profile accuracy was evaluated using PCC and MAE. PCC was computed as the Pearson correlation between the mean generated profile and the mean real target profile:

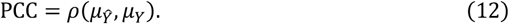

MAE measured the average absolute difference between these two mean profiles:

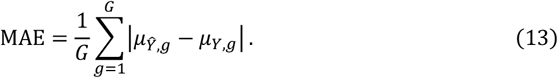

where *G* is the number of genes.

Distribution-level discrepancies were measured using maximum mean discrepancy (MMD) and E-distance. MMD was computed with an RBF kernel,

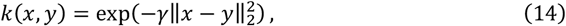

with 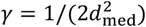 where *d*_med_ is the median pairwise Euclidean distance among real samples, with a small numerical stabilizer. The empirical squared MMD between generated and real samples was

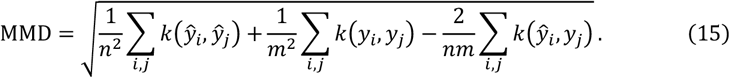

E-distance followed the energy-distance formulation:

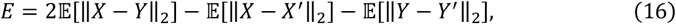

where *X, X*′ are independent generated samples and *Y, Y*′ are independent real samples.

### Randomization and statistical reporting

Randomness in the benchmark arose from train-test splitting, random pairing of unpaired single-cell profiles, sampling of target-domain control cells, model initialization, minibatch ordering and stochastic generation. All random procedures used fixed seeds specified in the released benchmark configurations. Unless otherwise noted, repeated experiments were run three times with independent training and evaluation. For published baselines with a fixed trained model and stochastic generation, we repeated only the generation step three times.

Results are reported as mean±s.d. across repeated runs or held-out evaluation units, as indicated in each figure. Standard deviations were computed within comparable replicate settings and were not pooled across distinct cell types, species, compounds or time points.

## Supplementary Figures

**Supplementary Figure 1.**
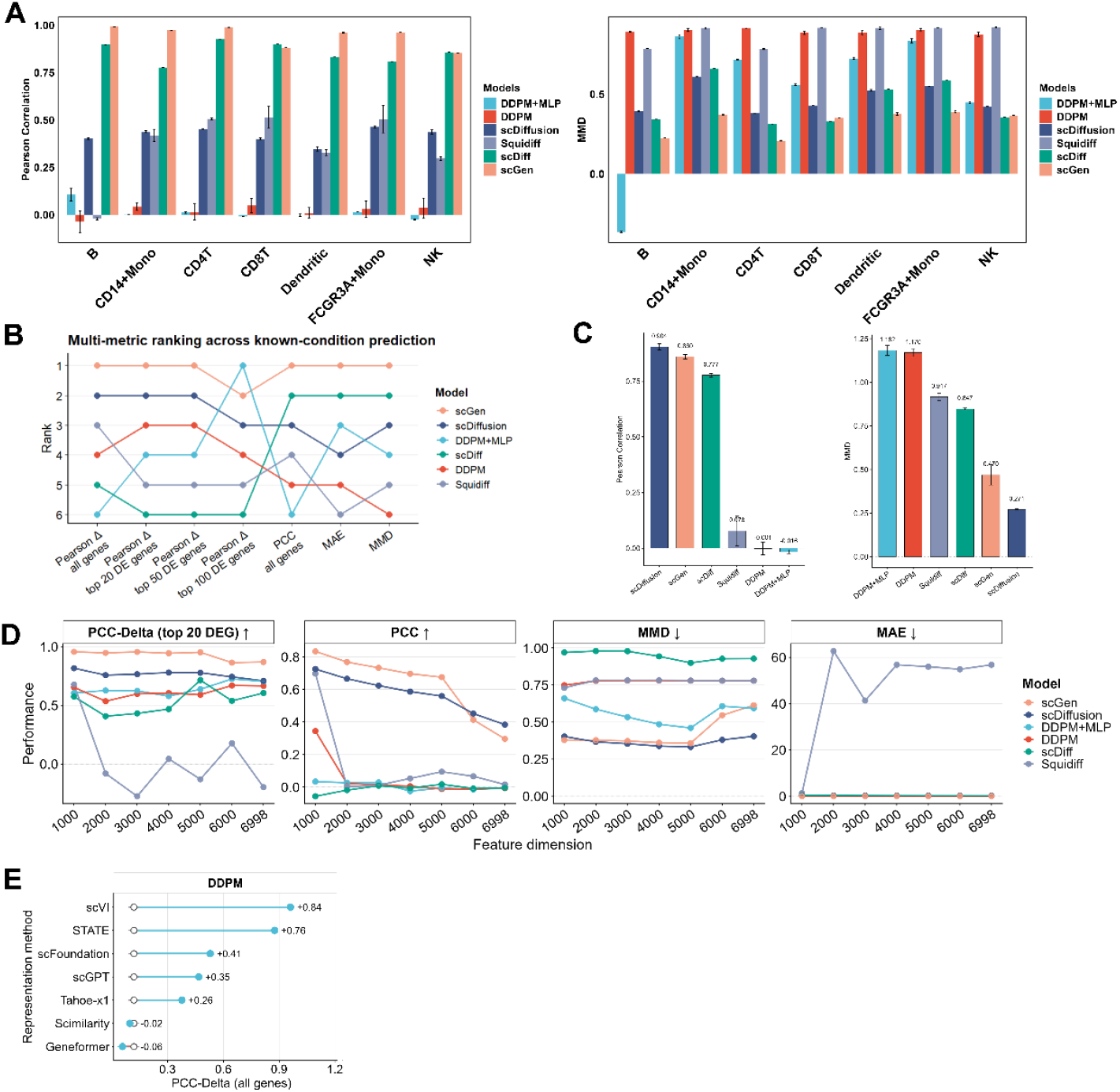
Additional evaluation of known-condition perturbation response prediction. (A) Model performance across seven cell types in the Kang PBMC dataset under the known-condition setting. Performance was evaluated using Pearson correlation and MMD for each cell type. (B) Multi-metric ranking of models in the single-cell known-condition benchmark. Models were ranked across gene-level correlation, DEG-based correlation, PCC, MAE and MMD metrics. Lower ranks indicate better performance. (C) Bulk RNA-seq perturbation response prediction for aflatoxin-treated samples. Models were evaluated using Pearson correlation and MMD under the known-condition setting. (D) Effect of input feature dimensionality on model performance in the CD4 T-cell known-condition task. Models were evaluated across different numbers of input genes using PCC-Delta for the top 20 differentially expressed genes, PCC, MMD and MAE. (E) Effect of representation choice on DDPM performance in the CD4 T-cell known-condition task. DDPM was combined with different representation models, and performance was evaluated using PCC-Delta across all genes.

**Supplementary Figure 2.**
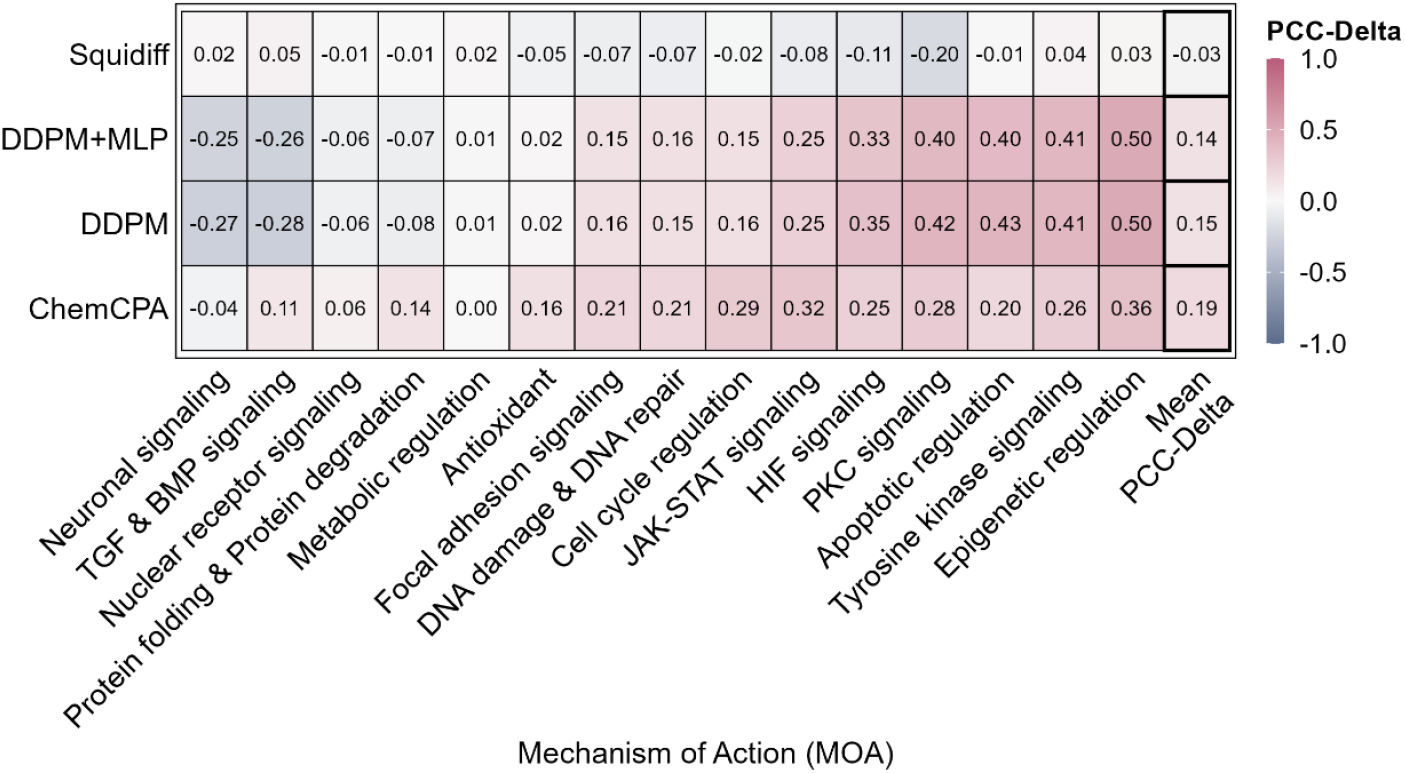
Within-MOA unseen drug prediction. Models were trained and tested within each MOA using predefined drug-level splits. Values indicate PCC-Delta for held-out drugs, and the final column shows the mean across MOAs. Higher values indicate better prediction performance.

**Supplementary Figure 3.**
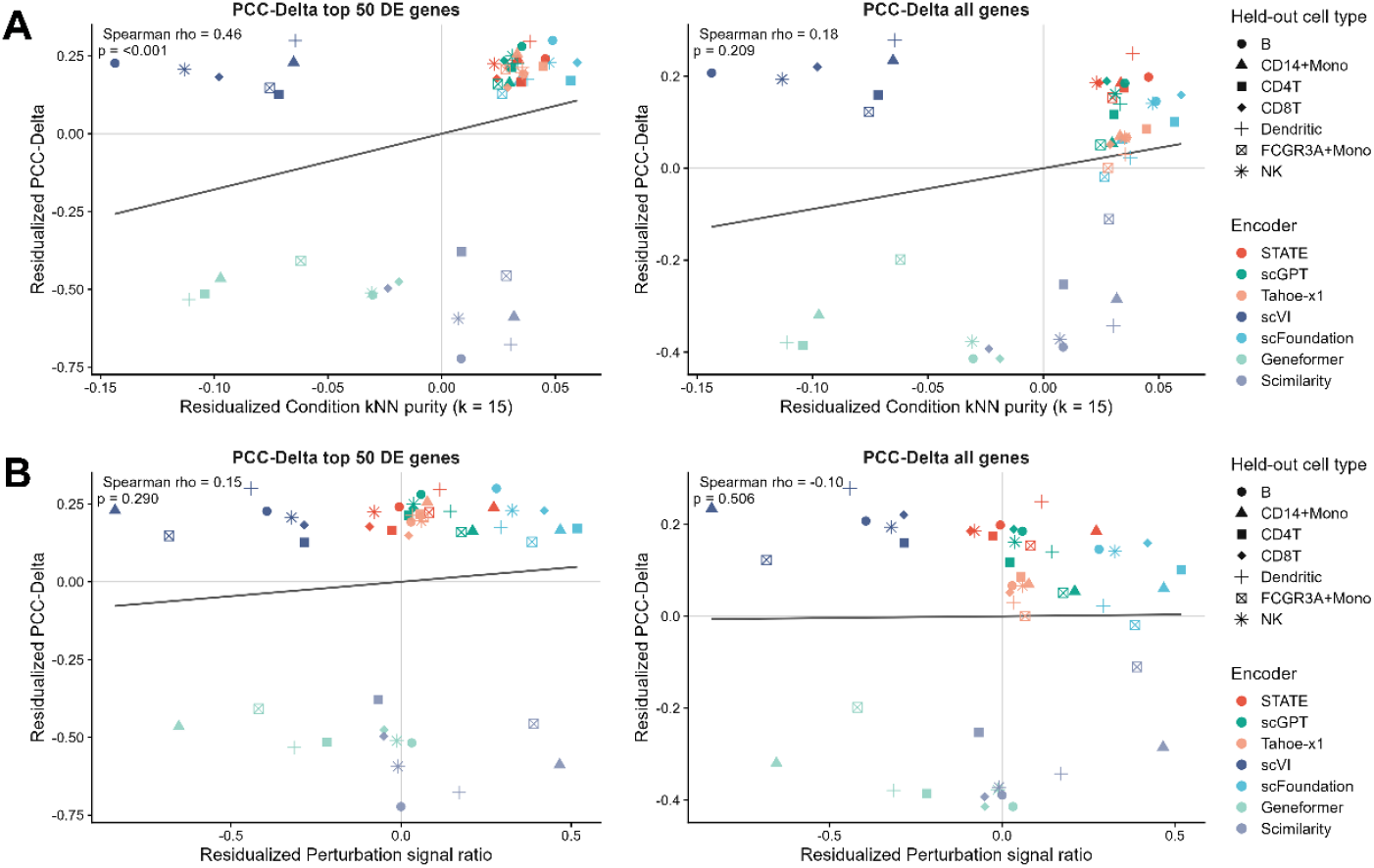
Perturbation-aware embedding structure in the unseen-cell-type task. (A) Association between condition k-nearest-neighbor (kNN) purity and DDPM prediction performance. Condition kNN purity was calculated as the fraction of the 15 nearest neighbors with the same perturbation condition, Control or IFN perturbed, in the encoder embedding space. (B) Association between perturbation signal ratio and DDPM prediction performance. Perturbation signal ratio was defined as the IFN perturbed–Control centroid distance normalized by within-condition variation. To account for differences in prediction difficulty across held-out cell types, both representation metrics and PCC-Delta values were centered within each held-out cell type before correlation analysis. DDPM performance was evaluated using PCC-Delta on the top 50 differentially expressed genes and all genes.

**Supplementary Figure 4.**
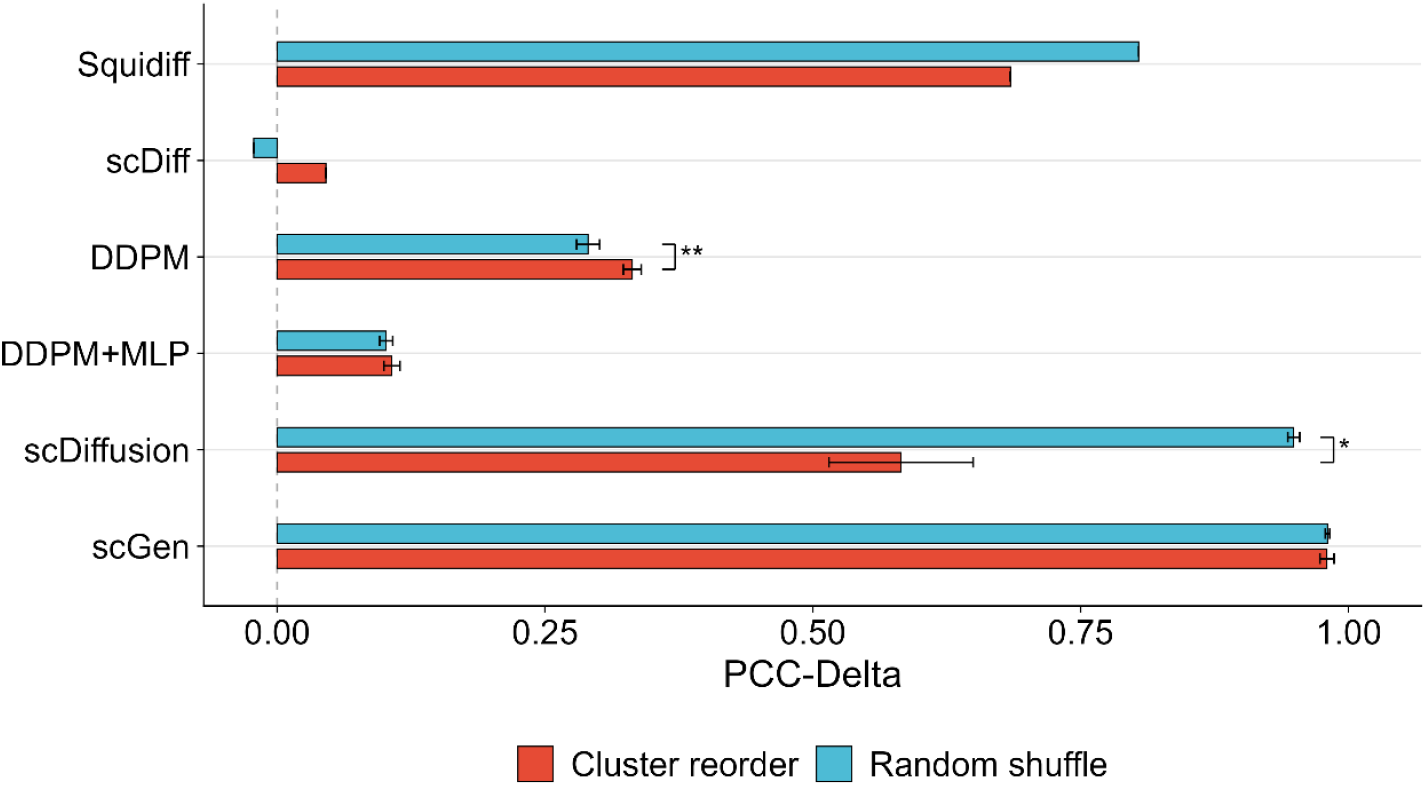
Gene-order perturbation in CD4 T-cell known-condition prediction. PCC-Delta is shown for six models under two input-order settings: clustering-based gene ordering and random gene shuffling. Both settings used the same 1,000 highly variable genes, and the assigned gene order was kept fixed across training and validation data. Bars show the mean across three runs, and error bars denote s.d. Within each model, the two ordering strategies were compared using two-sided Welch’s t-tests based on summary statistics (n = 3 per setting). Significant differences are indicated as *P < 0.05 and **P < 0.01.

## Supplementary Tables

**Supplementary Table 1.**
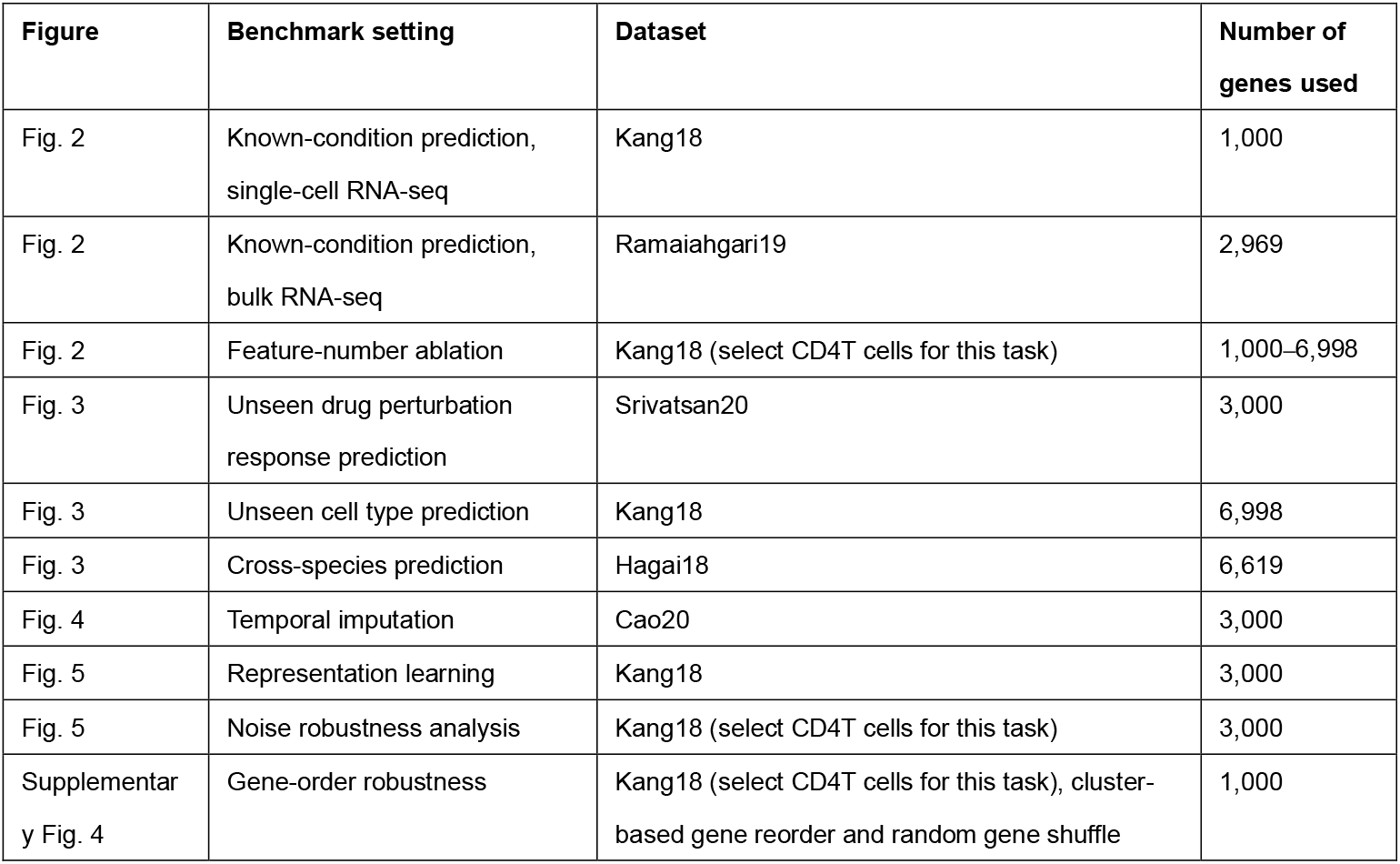
Dataset used in each benchmark task.

## Notes

https://github.com/ZijunSong/PertDiffBench

### Competing Interest Statement

The authors have declared no competing interest.

